# Null models for the Opportunity for Selection

**DOI:** 10.1101/2022.06.30.498333

**Authors:** R. S. Waples, Thomas E. Reed

## Abstract

Crow’s ‘Opportunity for Selection’ (*I*=variance in relative fitness) is an important albeit controversial eco-evolutionary concept, particularly regarding the most appropriate null model(s). Here we treat this topic in a comprehensive way by considering opportunities for both fertility selection (*I_f_*) and viability selection (*I_m_*) for discrete generations, both seasonal and lifetime reproductive success in age-structured species, and for experimental designs that include either a full or partial life cycle, with complete enumeration or random subsampling. For each scenario, a null model that includes random demographic stochasticity can be constructed that follows Crow’s initial formulation that *I*=*I_f_*+*I_m_*. The two components of *I* are qualitatively different. Whereas an adjusted *I_f_* (*Δ_If_*) can be computed that accounts for random demographic stochasticity in offspring number, *I_m_* cannot be similarly adjusted in the absence of data on phenotypic traits under viability selection. Including as potential parents some individuals that die before reproductive age produces an overall, zero-inflated-Poisson null model. It is always important to remember that (1) Crow’s *I* represents only the opportunity for selection and not selection itself, and (2) the species’ biology can lead to random stochasticity in offspring number that is either overdispersed or underdispersed compared to the Poisson (Wright-Fisher) expectation.

## INTRODUCTION

In 1958, James Crow introduced what he termed the ‘Index of Total Selection’ (*I*), which subsequent authors generally refer to as the ‘Opportunity for Selection’ (Arnold and Wade 1984; Clutton-Brock 1988). *I* is the variance in relative fitness (Walsh and Lynch 2018), and it sets an upper limit to the rate of evolutionary change by selection. Over the years, Crow’s index has generated a good deal of interest (Wade and Arnold 1980; Brodie at al. 1995; Ruzzante et al. 1996), as well as its share of controversy (Downhower et al. 1987; Fairbairn and Wilby 2001; Jennions et al. 2012). When Crow’s index is based on the most direct measure of fitness (the number of offspring, *k*, produced by each potential parent), it is calculated as the squared coefficient of variation in offspring number:

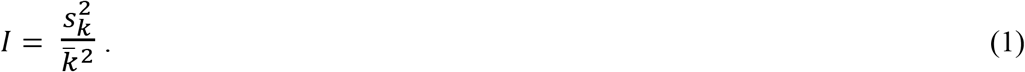

One unfortunate characteristic of *I* calculated as in Equation 1 is that the result is sensitive to 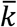, which means that conclusions about the Opportunity for Selection can depend as much or more on aspects of experimental design (sampling effort; life stage at which offspring are sampled) as they do on the biology of the focal species. Because an uneven sex ratio creates different mean offspring numbers for male and female parents, this dependence on 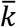 also complicates comparisons of the Opportunity for Selection between sexes within the same population.

Waples (2020) suggested a simple solution to the dependence of *I* on mean offspring number. Under the Wright-Fisher model of random reproductive success, the variance in offspring number is binomial (closely approximated by the Poisson variance), which means that 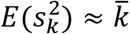. It follows that the random expectation for *I* under Wright-Fisher reproduction is 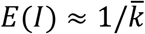. This latter result had been pointed out by others (e.g., Downhower et al. 1987), but Waples (2020) showed that subtracting this random expectation from the raw *I* produces an adjusted index 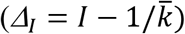 that is independent of 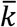. Use of *Δ_I_* rather than raw *I* thus can potentially facilitate comparisons of the Opportunity for Selection across species and across studies; within a study, it can also facilitate comparisons across different replicates (e.g., multiple seasonal episodes of reproduction) and between the sexes. Because this adjustment involves subtracting the expected contribution to *I* from random demographic stochasticity, it also can serve as a null model for the Opportunity for Selection, and null models increasingly play an important role in both ecology and evolutionary biology (Harvey et al. 1983; Maddison and Slatkin 1991; Gotelli and Ulrich 2012; Steiner and Tuljapurkar 2012; Farine 2017; van Daalen and Caswell 2017; Ross et al. 2020).

However, the *Δ_I_* approach described by Waples (2020) has some important limitations. First, the theory behind *Δ_I_* was developed using a generalized Wright-Fisher model that assumed discrete generations, and the numerical evaluations performed by Waples (2020) also modeled discrete generations. Most real species, however, are age structured. Although Waples (2020) discussed some of the possible implications of age structure for *Δ_I_*, it is important to provide a more rigorous treatment for this key topic. Second, in many terrestrial vertebrates, physiological limits on litter or clutch size constrain the variance in annual offspring number, leading to underdispersion that makes the Poisson model a poor description of random demographic stochasticity. The third major limitation is that the *Δ_I_* approach followed the Wright Fisher model in assuming that all potential parents had an equal opportunity to produce offspring. The order of events in the null model was therefore reproduction, followed by random survival until the stage at which offspring are sampled. In contrast, Crow’s formulation of *I* used a more general definition of fitness to include variation in survival as well as variation in offspring number.

A comprehensive evaluation of null models for the Opportunity for Selection therefore must consider several additional scenarios beyond the one considered by Waples (2020): (1) explicitly accounting for age structure; (2) enumerating potential parents at an early life stage, followed by mortality prior to maturity and eventual reproduction by the survivors; (3) ignoring reproduction and only considering survival between two life stages; and (4) evaluating factors that cause random demographic stochasticity in annual reproductive success to deviate from the Poisson expectation. Our goal here is to provide this comprehensive evaluation of null models for Crow’s index. Analytical results are obtained for three different life-history scenarios: discrete generations; seasonal reproduction in age-structured species; and lifetime reproductive success for iteroparous species. For each life-history scenario, we consider a range of experimental designs that lead to different, ordered combinations of survival and/or reproduction at different life stages. Simulations are used to confirm analytical results and to illustrate practical applications for a range of experimental designs.

## METHODS

### Notation and Experimental Design

The core data under consideration here are means and variances in the number of offspring (*k*) per potential parent. Collecting these data requires taking samples of individuals in two ordered time periods, with *T*_2_ later in time than *T*_1_. At *T*_1_, a set of *N* potential parents is identified, and the first sample of individuals is randomly collected from this set. For simplicity, in the treatments below it is assumed that the potential parents are exhaustively sampled, but randomly subsampling potential parents does not qualitatively change the results. At *T*_2_, a random sample of size *N_Off_* is collected from the offspring arising from the potential parents sampled in *T*_1_. The sample mean number of offspring per parent is therefore 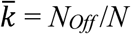, whereas the sample variance 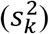 is a random variable that depends on various factors as described below. In general, it is best to measure reproductive success across a full life cycle (so at the same age or stage (*T*_2_ = *T*_1_), but one life cycle later); however, in many real-world applications this is difficult or impossible to accomplish, hence the more general treatment here.

The above applies generally to discrete generations or to seasonal reproduction in age-structured species. For evaluation of lifetime reproductive success (*LRS*) in iteroparous species, it is necessary to integrate information about offspring number across parental lifespans. We use the “•” to denote metrics (such as 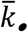 and 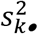) associated with lifetime reproduction. Variance in *LRS* can be affected by temporal autocorrelations of reproductive output over time (‘persistent individual differences’; Lee et al. 2011) and by positive or negative correlations between reproductive success and survival (as commonly occur, for example, with intermittent breeding). The null model we use assumes these correlations vary randomly around 0, but researchers should keep potential effects on *LRS* in mind.

### Components of the Opportunity for Selection

Crow (1958) defined *I* in terms of the mean and variance in offspring number among a group of individuals enumerated at birth. Subsequent mortality divides this initial cohort into two subgroups: those that survive to reproductive age (fraction *v*) and those that do not (fraction 1-*v*). This stratification of the cohort into winners and losers *vis a vis* premature mortality allowed Crow to identify two components of *I*: one due to mortality (*I_m_*) and one due to differential fertility (*I_f_*). Across the entire life cycle, overall *I* = *I_m_* + *I_f_*, provided survival and reproduction are independent (see Arnold and Wade 1984). Whereas *I_f_* can be a complicated function of the distribution of offspring number among surviving parents, the component due to mortality is simple and straightforward: *I_m_* = (1-*v*)/*v* = the fraction of potential parents that die before reaching sexual maturity, divided by the fraction that survive (Crow 1958).

### Discrete Generations

With discrete generations, each birth cohort represents a generation, and all members simultaneously progress through the various life stages (*J*_1_, *J*_1_,… *J_n_*, *A*), where the *J* represents various juvenile stages (*J*_1_ = zygotes) and *A* is the single adult stage; Figure 1. The cycle is continuous, without any official start or end. The most general model involves inventorying individuals at two ordered time points: *T*_1_ = any of the above stages, and *T*_2_ = some subsequent stage. We restrict analysis to scenarios in which the maximum time frame is one full generation, but we also allow for consideration of analyses that cover only part of a generation. In this model, four general scenarios are possible, depending on the experimental design: I) *T*_1_ and *T*_2_ encompass only survival and not reproduction (e.g., *T*_1_ = *J*_2_ and *T*_2_ = *J*_n>2_); II) *T*_1_ and *T*_2_ encompass only reproduction and not survival (*T*_1_ = *A* and *T*_2_ = *J*_1_); III) *T*_1_ and *T*_2_ encompass survival followed by reproduction (e.g., *T*_1_ = *J*_3_ and *T*_2_ = *J*_2_, as shown in Figure 1); IV) *T*_1_ and *T*_2_ encompass reproduction followed by survival (e.g., *T*_1_ = *A* and *T*_2_ = *J*_2_). In Results we consider appropriate null models for each of these scenarios.

**Figure 1.**
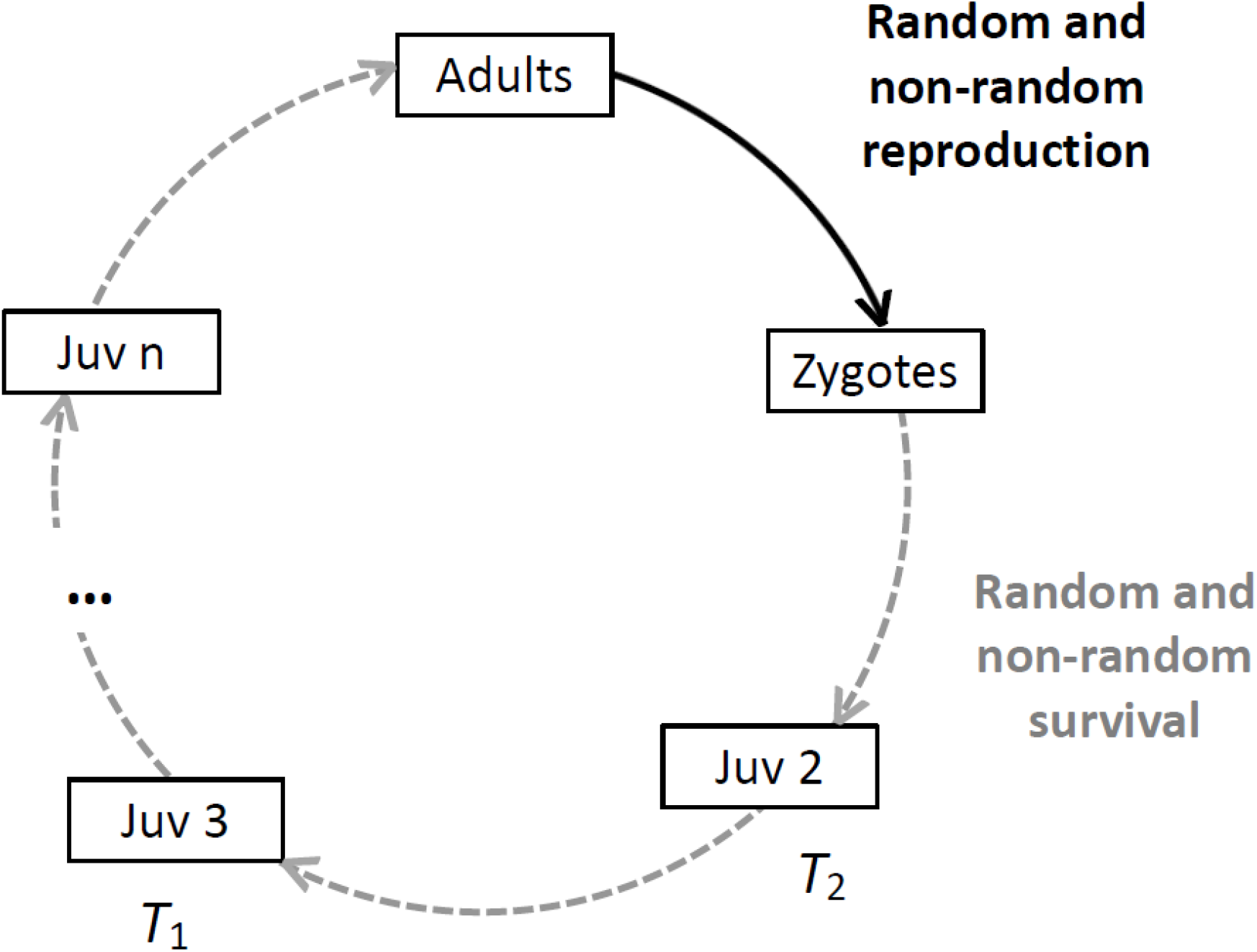
Generic life cycle model for a species with discrete generations. Several life stages can be identified, but at any given time all individuals are in the same life stage. Reproduction only occurs at the adult stage (black), whereas earlier life stage (gray) experience sequential episodes of mortality. *T*_1_ is the life stage at which potential adults are sampled, and *T*_2_ is the life stage at which their offspring are sampled.

### Overlapping Generations

With overlapping generations, individuals of different ages live and reproduce at the same time, and some individuals survive to reproduce in subsequent time periods. The model used here is discrete-time, with age indexed by *x*, and assumes that reproduction is concentrated within seasonal time periods, hereafter without loss of generality assumed to be years. This corresponds to the birth-pulse model of Caswell (2001), which is applicable to a wide range of taxa. Some key features of evolutionary demography for populations like this can be summarized by specifying the age-specific vital rate *v_x_* = probability of surviving from age *x* to age *x*+1. [A typical life table also specifies age-specific fecundity, but under our null model fecundity is assumed to be constant.] The cumulative survivorship function *l_x_* incorporates information about age-specific survival across individual lifespans and hence determines population age structure (Table 1). Some key ages can be identified: 0 = newborns, which is the closest equivalent to zygotes; *α* = age at maturity (assumed to be fixed); and *ω* = maximum age. It is also important to define the set of individuals for which the mean and variance in offspring number will be computed. We define *z* as the age at enumeration of potential parents and define the set of potential parents as individuals with age ≥ *z*, where *z* is in the range (0,*α*). [Technically, setting α<*z*≤*ω* is possible but is not considered here, as then reproductive success would be assessed for only a subset of mature individuals.]

**Table 1.** Demographic data for the age-structured population illustrated in Figure 2, which has 10 age classes (0-9), constant survival at the rate *v* = 0.7/season, a fixed cohort size of *N_0_* = 800 newborns, and reaches sexual maturity at age α = 3. Data apply to a single sex and could vary by sex. *l_x_* is cumulative survivorship through age *x*; *N_x_* = *N_0_l_x_* = number of individuals alive at age *x*; *D_x_* = number of individuals that die between ages *x* and *x*+1; *q* = age at death, scaled to reflect the number of years of the adult lifespan during which the individual potentially could have reproduced (ages α:ω = 3:9 in this example); *D_q_* = *D_x_* for the years of the adult lifespan, re-indexed by *q*. In the example in Figure 2, the age at enumeration (which defines the set of potential parents for which reproductive success will be assessed) is *z* = 1. For analysis of annual reproduction (Figure 3), 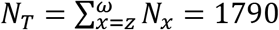 and the total population of adults is 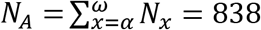. These same data can be used for the analysis of lifetime reproductive success (Figure 5), which focuses on the number of individuals in a cohort that reach age at enumeration (*N_z_* = 560) or age at maturity (*N_α_* = 274).

| Age | $l_x$ | $N_x$ | $D_x$ | Age at<br>Death ($q$) | $D_q$ |
| --- | --- | --- | --- | --- | --- |
| 0 | 1 | 800 | 240 |  |  |
| 1 | 0.7000 | 560 | 168 |  |  |
| 2 | 0.4900 | 392 | 118 |  |  |
| 3 | 0.3430 | 274 | 82 | 1 | 82 |
| 4 | 0.2401 | 192 | 58 | 2 | 58 |
| 5 | 0.1681 | 134 | 40 | 3 | 40 |
| 6 | 0.1176 | 94 | 28 | 4 | 28 |
| 7 | 0.0824 | 66 | 20 | 5 | 20 |
| 8 | 0.0576 | 46 | 14 | 6 | 14 |
| 9 | 0.0404 | 32 | 32 | 7 | 32 |
| $N_T$ | | 1790 | | | |
| $N_A$ | | 838 | | | |

For iteroparous species, reproductive skew and the Opportunity for Selection are commonly assessed in two different ways: 1) annual reproductive success across all individuals alive during a single season; and 2) lifetime reproductive success among individuals of the same birth cohort.

### Annual reproduction

With annual reproduction, we are interested in the mean and variance in offspring number for all individuals with age ≥ *z* that are alive at a given time. Assuming the population has stable age structure and a fixed cohort size of *N_0_* newborns, the vector of numbers alive at age is *N_x_* = *N_0_l_x_*, with *l_0_* = 1, which leads to Σ*N_x_* = *N_0_∑l_x_* (Table 1). Therefore, if reproductive success is assessed for all individuals age *z* and older, the total number of potential parents for which 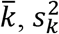, and *I* are computed is (see Table 1 and Figure 2)

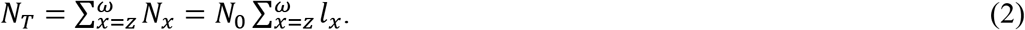

The age at which offspring are sampled also is a factor that can affect calculation of 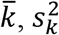, and *I*. However, the expectations for 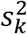 and *I* are conditional on the mean offspring number rather than age (Waples 2020), so it is sufficient to express these expectations in terms of 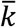.

**Figure 2.**
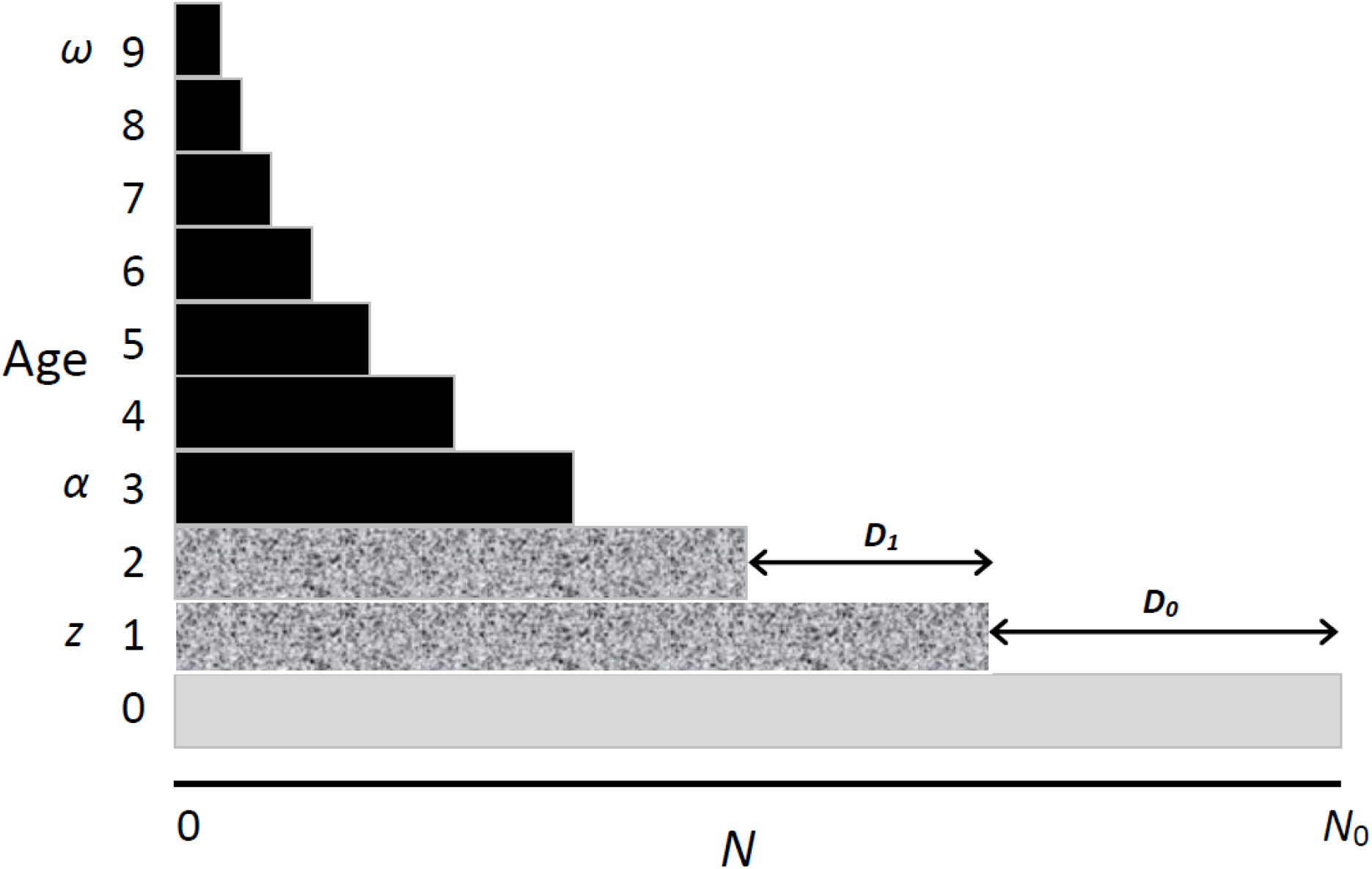
The stable-age-structure age pyramid for a hypothetical population with constant recruitment of *N_0_* newborns per year, which subsequently survive to the next age with fixed probability *v* = 0.7, as in Table 1. Solid arrows and *D_x_* values indicate the number of individuals that die after reaching age *x* but before reaching age *x*+1. Maximum age is ω = 9 and age at maturity is α = 3, so the black rectangles represent all adults (*N_A_*) alive at any given time. Annual reproductive success (see Figure 3) is assessed for all *N_T_* individuals with age ≥ age at enumeration, which in this example is *z*=1. Individuals aged 1 and 2 (shaded rectangles) are not sexually mature and hence all produce 0 offspring in the current season. Analysis of lifetime reproductive success (see Figure 5) focuses on individuals in a single birth cohort. The fraction of immature individuals in a cohort that are expected to die before reaching sexual maturity is a simple function of their current age and survival rate, *v*.

### Lifetime reproduction

In contrast to annual reproductive success, which reflects offspring produced in a single time period by mixed-age individuals, *LRS* is typically assessed for a single birth cohort of individuals by integrating their annual reproductive success across entire lifespans. The null model for *LRS* is more complicated because, in addition to accounting for random reproduction within years, it also has to account for the fact that some individuals live longer than others and thus have more opportunities to accumulate offspring. The maximum number of years or reproductive seasons in which any individual can reproduce is the adult lifespan: *AL* = ω – α + 1. In addition, mean and variance of *LRS* are also affected by the age at which the cohort is defined; if the cohort includes immature individuals, this has to be accounted for by a term for premature mortality, as is the case for discrete generations and annual reproduction. Finally, variance in *LRS* can be affected by correlations between survival and reproduction, or between reproduction by the same individual in different years/seasons. In the null model, these correlations are assumed to have expectations = 0.

Let a cohort be defined by all individuals that survive to age *z*. Then, the number of individuals for which the mean and variance in lifetime offspring number are computed is *N_z_* = *N_0_l_z_*. Using vital rates from a standard life table, it possible to compute the expected value of 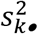 by grouping individuals by age at death (Waples et al. 2011; Waples 2022). These ages at death are equivalent to treatments in an ANOVA analysis; assuming a fixed age at maturity, individuals that die at the same age have the same number of annual opportunities to reproduce and, under the null model, the same *E*(*LRS*).

### Simulations

To verify the accuracy of the analytical expectations, we conducted computer simulations of random survival and reproduction using the null models described here. Population demography followed Table 1, and computer code is available in Supporting Information. Each replicate started with a fixed cohort size of 800 newborns, after which individuals survived to the next age with probability 0.7. This allowed for random variation in realized numbers-at-age (*N_x_*) in each replicate. In Figures 3 and 5, the expected results use the stable-population *N_x_* values shown in Table 1, whereas observed results are empirical medians across 1000 simulated replicates.

**Figure 3.**
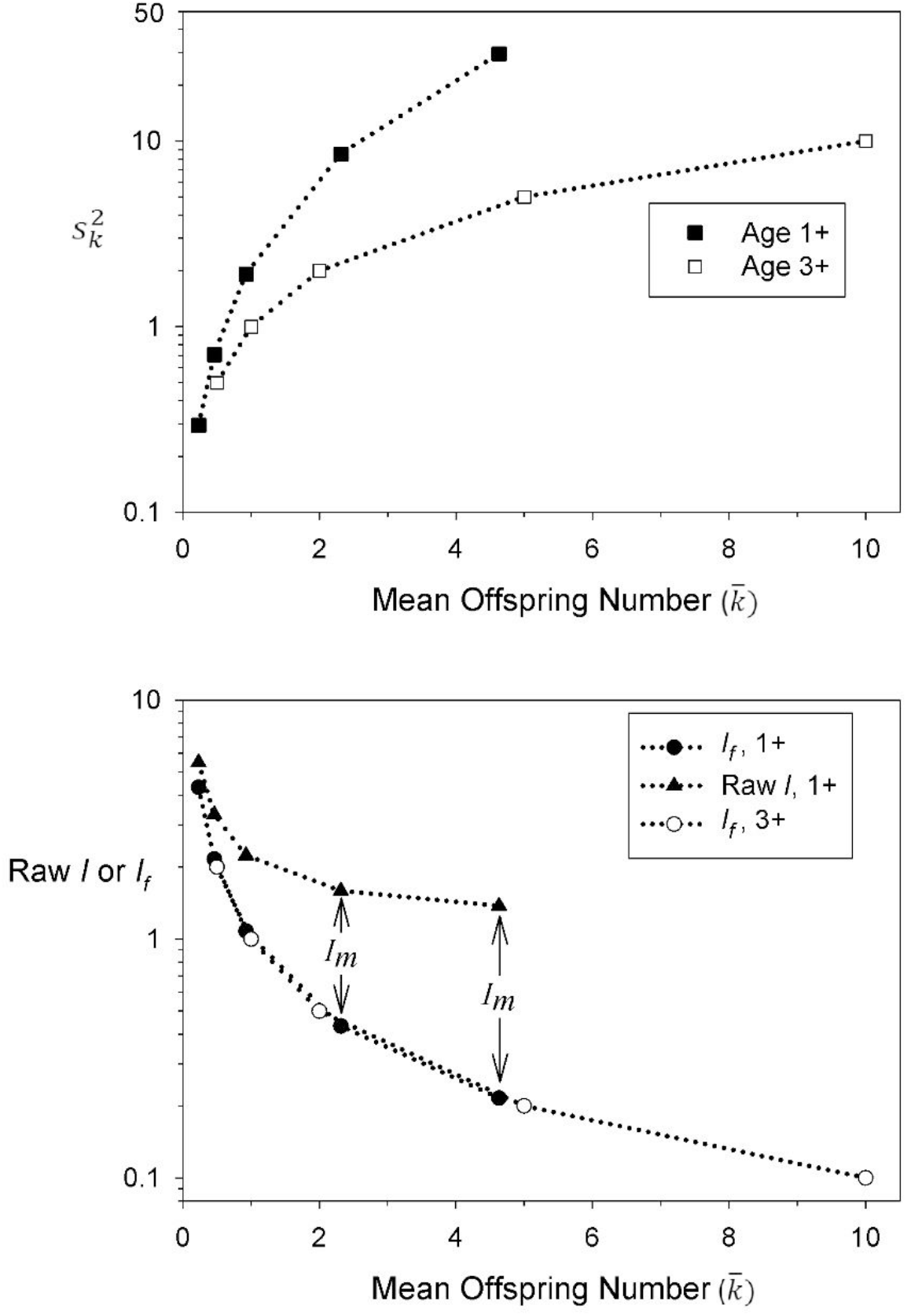
Results for annual reproductive success under the null model, for a population with age structure as shown in Table 1 and Figure 2. Dotted lines are expected results based on equations in the text; symbols are medians of 1000 replicate simulations. Note the log scales on the *Y* axes. Means and variances were calculated in two ways: 1) for all individuals with age ≥ age at enumeration (“1+”); and 2) only for individuals with age ≥ age at maturity (“3+”). Top: 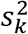 as a function of mean annual offspring number. The ratio of the number of age-1 individuals that die before reaching age 3 (286) to the number that survive (274) is > 1, so when all the age-1 individuals are included as potential parents (solid squares), 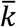 is reduced by a factor of more than 2, and the variance in offspring number is inflated by deterministic zeroes for those 286 individuals. Bottom: raw values of Crow’s *I* are also inflated by inclusion of immature individuals as potential parents (solid triangles). After subtracting the constant value of *I_m_* = 286/274 = 1.04 to account for the fraction of immature individuals, the net value of *I_f_* accounts for random variation in fertility and its expected value is 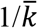 (solid circles), which is the same result obtained when reproductive success is assessed using only mature individuals (open circles).

**Figure 4.**
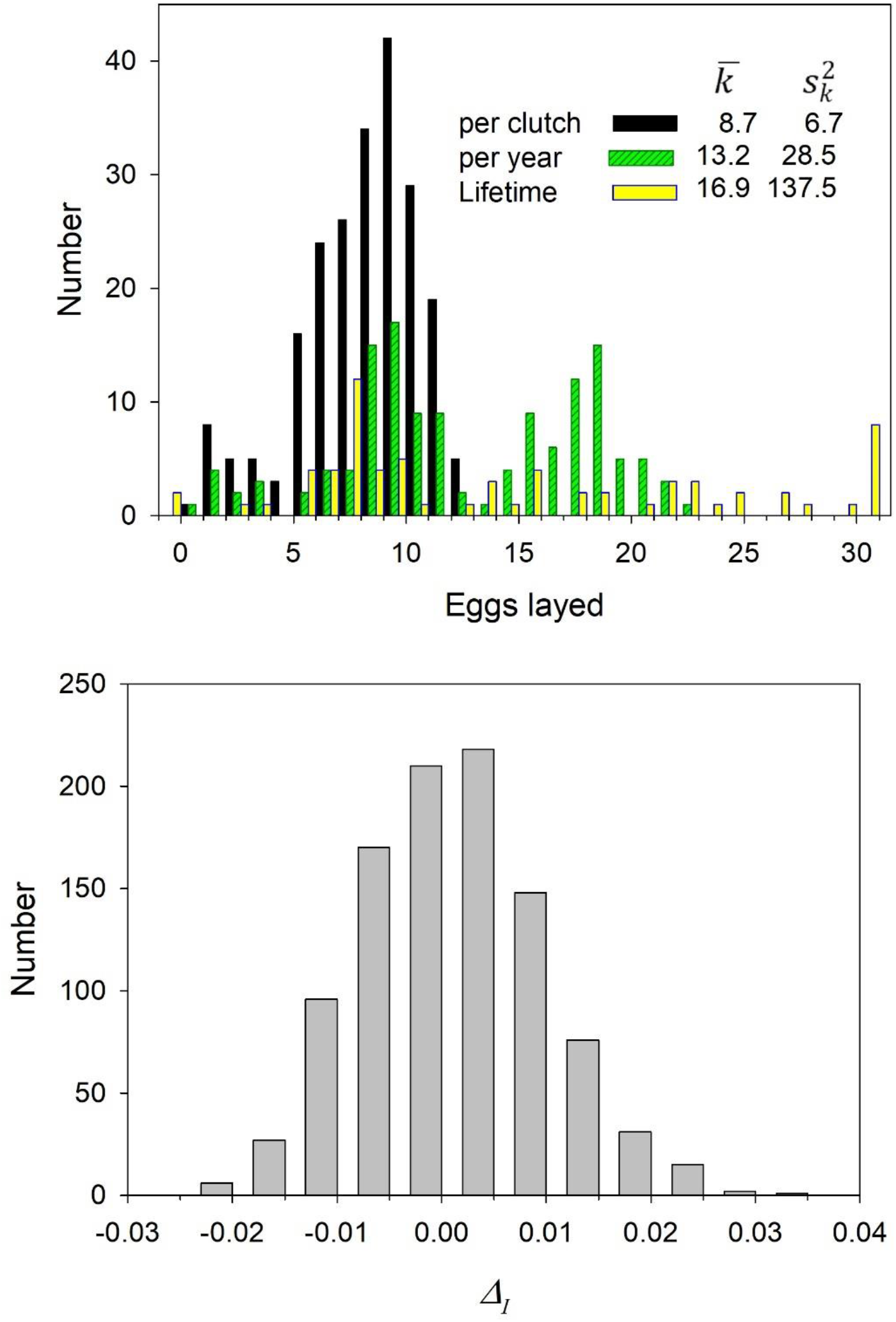
Distribution of egg production in the population of great tits from Hoge-Veluwe, the Netherlands, in 1980. Top: mean and variance in clutch size (black bars), total eggs per year (green bars), and total eggs per lifetime for the 1980 cohort (yellow bars). Bottom: distribution of the generalized *Δ_I_* across 1000 replicate populations simulated using the generalized Poisson distribution with parameters λ_1_ and λ_2_ matched to the empirical mean and variance of clutch size in the great tit.

**Figure 5.**
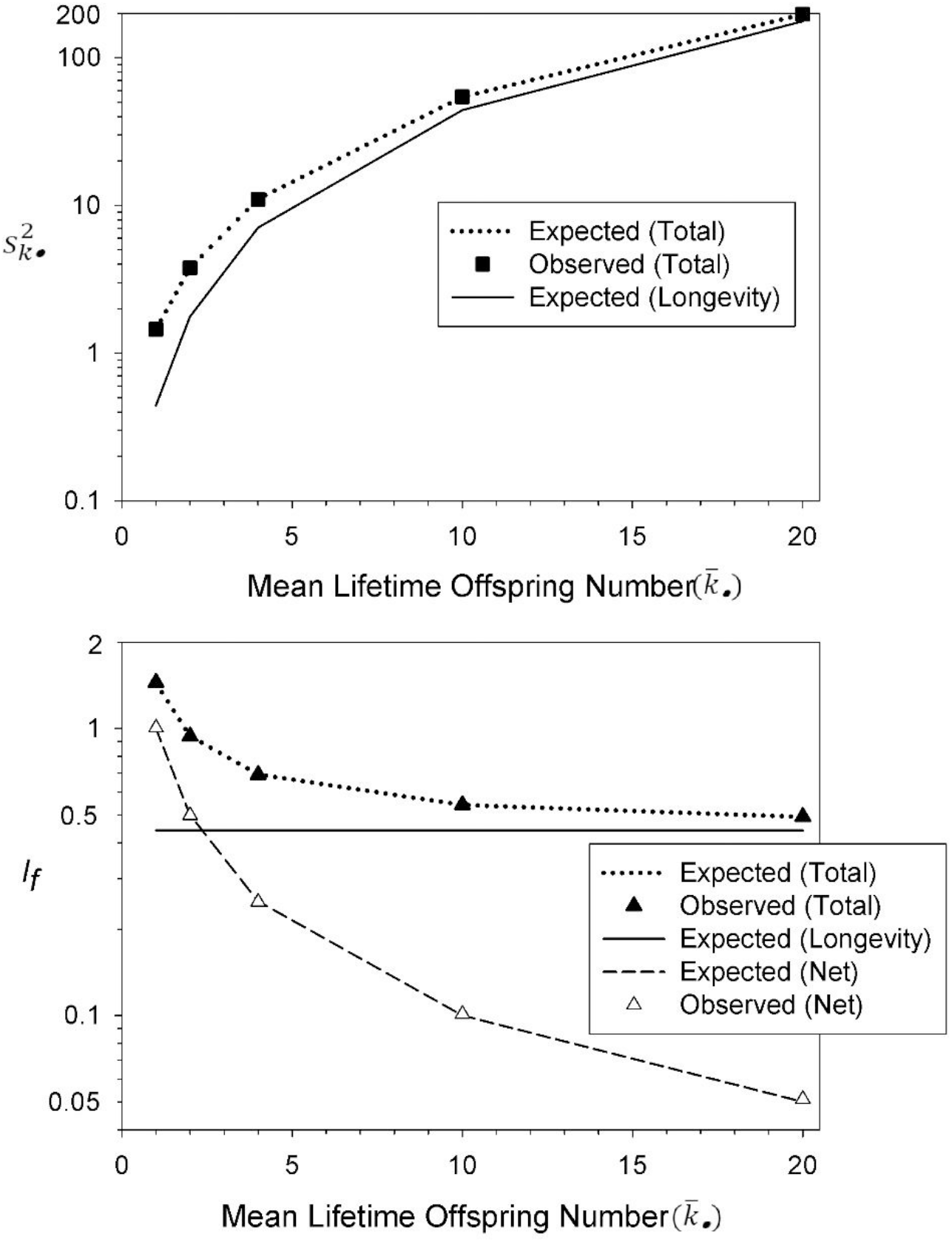
Results for *LRS* under the null model, using demographic data from Table 1. Lines are expected results based on equations in the text; symbols are medians of 1000 replicate simulations. In all analyses, means and variances were calculated only for the mature individuals in a cohort. Note the log scales on the *Y* axes. Top: variance in lifetime reproductive success as a function of 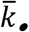. Dotted line and filled squares show the total variance; solid line shows the theoretical expectation for the component arising from random variation in longevity. Bottom: Crow’s Opportunity for Selection component *I_f_* as a function of 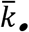. Dotted line and filled triangles show the total *I_f_*; solid line shows the theoretical expectation for random variation in longevity (which is independent of mean offspring number), and dashed line and open triangles show the net *I_f_*; after subtracting the longevity component. After this adjustment, the expected value of the net 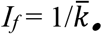.

## RESULTS

### Discrete Generations

#### Mortality only, no reproduction

This is the simplest scenario; as shown by Crow (1958), *I_m_* = (1-*v*)/*v*, where *v* is the fraction of individuals that survive to the later life stage and 1-*v* is the fraction that don’t. No reproduction is involved, so Δ_*I*_ as defined by Waples (2020) is not relevant here. Because all or none of this mortality could be random, there really is no quantitative null model here, in the same way there is for reproduction. With respect to mortality, a reasonable null model would be that in which individuals survive across a specific life stage or time period independent of any phenotypic traits. This null model then could be rejected by finding a significant covariance between survival and phenotype.

#### Reproduction only, no mortality

Conceptually, this is the first step in the Wright-Fisher model, since each individual contributes exactly the same number to the essentially infinitely-large pool of gametes, which then unite at random to form zygotes. In the null model, therefore, at this stage there is zero variance among individuals in offspring number. In the real world, all gamete and zygote pools are finite. Therefore, any actual analysis of empirical data that involves estimating mean and variance in offspring number per parent will of necessity involve either a) enumerating offspring at a later life stage than zygotes, or b) subsampling only some of the offspring, or c) both. These scenarios are described next.

#### Mortality followed by reproduction

Across a full life cycle, this scenario quantifies reproductive success of zygotes producing zygotes, and here we must consider both components of *I*. Crow defined *I_f_* in the familiar format as the sum of squared deviations of offspring numbers from the mean (in our notation, 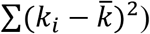. For the purposes of identifying a null model, a more convenient approach is to deal instead with sums of squared offspring numbers 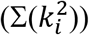.

We take advantage of the parametric definition of a variance as 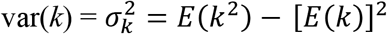. If we let *SS* = sum of squares = ∑(*k*^2^) and 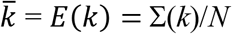 and ignore the (*N*-1) correction for sample variance, then this can be written as 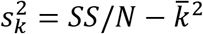. Rearrangement produces an expression for the total sum of squares: 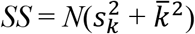. In the null model, individuals survive at random, and those that do survive are assumed to have a Poisson distribution of reproductive success, regardless at what stage the offspring are counted (so for the mature individuals, 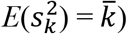).

If potential parents are sampled at some pre-adult life stage, and only the fraction *v* survive to age at maturity, at which point *N* remain, then the total number of individuals for which offspring numbers are counted is *N/v*. Let 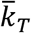 = overall mean offspring number for these *N/v* individuals. There are two groups: *N/v* – *N* do not survive to maturity and produce 0 offspring, and *N* do survive and produce on average 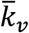 offspring each. The total number of offspring examined is 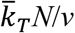, so 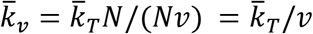. So the *N* individuals that survive to maturity produce a mean 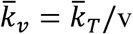 offspring each, and under the null model this is also = 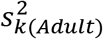. For the *N* adults, 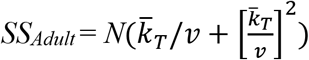, and this is also the total sum of squares, since those that died left no offspring. The total variance in reproductive success across the *N/v* individuals is

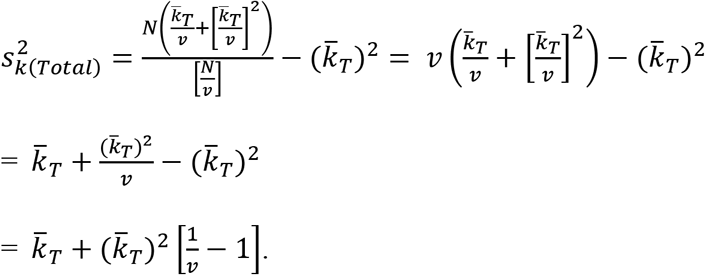

With *v* = 1 (no mortality prior to sexual maturity), this simplifies to 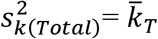, leading to 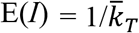, consistent with Δ_*I*_. More generally, under the null model,

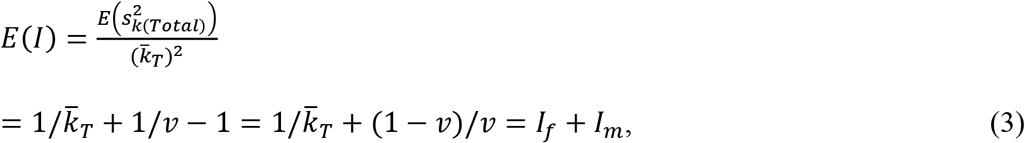

where *l_m_* = (1 – *v*)/*v* as defined by Crow (1958). This same result can be obtained using formulas for the mean and variance of a zero-inflated Poisson distribution (see Supporting Information). As the fraction that survive (*v*) declines, the term (1 – *v*)/*v* becomes arbitrarily large, and that is what inflates the Opportunity for Selection compared to the Poisson expectation that only applies to reproduction. Note that *I_f_* in Equation 3 corresponds conceptually to the opportunity for fecundity selection via *zygote* fertility (i.e., how many zygotes each newly-conceived individual produces), not adult fertility.

#### Reproduction followed by mortality

When reproductive success is measured per adult, a qualitative difference occurs in the Opportunity for Selection because there is no *l_m_* component, which as defined by Crow (1958) applies only to pre-reproductive mortality. Instead, any mortality that occurs before offspring are enumerated is folded into the *I_f_* component.

Here, Wright-Fisher is an appropriate null model: no drift occurs at the first step (reproduction), so all variance in offspring number arises from random mortality between the (conceptually infinite) zygote stage and the stage at which offspring are sampled. The following two scenarios produce identical results from a statistical perspective: (1) random sampling a later life stage that has experienced random mortality, and (2) randomly sampling from the earlier life stage. Under Wright-Fisher, 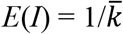, regardless whether the sampled individuals represent the entire population of offspring alive at the stage sampled, or just a random subset. This is the scenario Δ_*I*_ was designed to deal with. If offspring mortality is family-correlated (such that offspring from different families have different probabilities of surviving), that will increase the variance in offspring number (Crow and Morton 1955) and increase the Opportunity for Selection above the Poisson expectation. In that scenario, Δ_*I*_ will on average be positive, even when all individuals have an equal expectation of reproductive success. However, by itself a positive value of Δ_*I*_ would not indicate whether the departure from Wright-Fisher dynamics is due only to non-random reproduction by the parents, only to non-random survival of offspring, or both.

#### Annual Reproduction

In this case, means and variances of offspring number are computed across all or a subset of individuals that are alive in a given year or season. Here the null model also involves two groups: those that are mature and those that are not, and we want to determine *E*(*I*) for all *N_T_* individuals with age ≥*z*. Let *N_A_* = number of adults of all ages, let 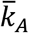 = mean offspring number for adults and let 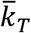 = mean offspring number for all *N_T_* individuals. The total number of offspring examined is thus 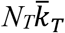, and these are all produced by the *N_A_* adults, so 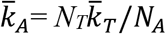, and under the null model this is also 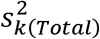 for adults. So for the *N_A_* adults,

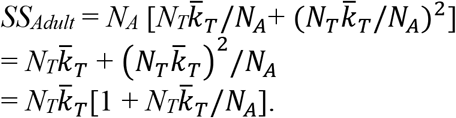

*SS_Adult_* is also the total *SS_T_* for all *N_T_* individuals, so it follows that the total variance in offspring number is

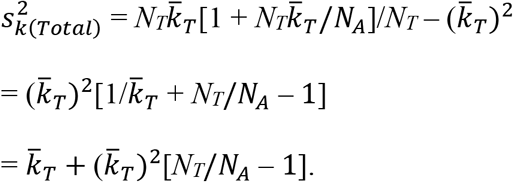

Therefore, 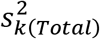 exceeds the random (Poisson) expectation 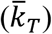 to the extent that *N_T_*>*N_A_*, and this effect can be attributed to including a subset of immature individuals with zero probability of producing offspring in the calculations of 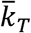 and 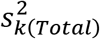. It follows that under the null model for annual reproduction,

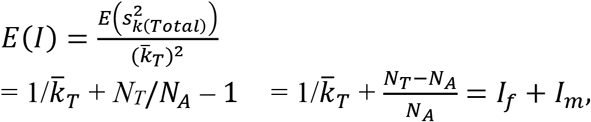

exactly as obtained under the discrete generation model (Equation 3). Note that in the term 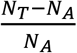, the denominator is the subset of the initial population that survives to age at reproduction and the numerator is the subset that do not, so this term is identical to *I_m_* as defined by Crow (1958). If only adults are included in the computation, *N_T_/N_A_* = 1 and 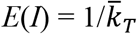, as is the case for discrete generations.

In the simulations of annual reproduction, observed values of 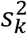 and Crow’s *I* agreed almost exactly with these expectations (Figure 3). After subtracting the component *I_m_* to account for premature mortality, the component of Crow’s index related to fertility (*I_f_*) is just the inverse of the mean offspring number, as proposed by Waples (2020).

The key ratio *N_T_/N_A_* can be expressed in terms of cumulative survivorship from age *z* to age α:

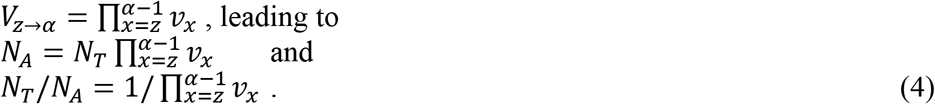

For the special case where annual survival rate is constant for the ages involved (all *v_x_* = *v*), *V*_*z*→*α*_ = *v^y^*, where *y* = α-*z* = the difference between the ages at maturity and enumeration, which is also the number of annual episodes of random mortality between the two ages. Equation 4 then simplifies to *N_T_/N_A_* = 1/*v^y^*. If *y* = 0, this reduces to *N_T_/N_A_* = 1, as expected.

Although age-specific vital rates like those in Table 1 are commonly reported for iteroparous species, many semelparous species do not have a fixed age at maturity [for example, Pacific salmon (Waples 2006), annual plants with seed banks (Nunney 2002), and monocarpic perennials (Vitalis et al. 2004)], and these species therefore have an interesting combination of traits normally associated with both discrete and overlapping generations. In these species, annual reproduction involves potential parents of mixed ages, just as it does for iteroparous species. For semelparous species with variable age at maturity, therefore, annual reproduction can be analyzed using the same framework described in this section. The difference is that for semelparous species this annual reproduction also represents their lifetime reproductive output; for iteroparous species, *LRS* can be analyzed as described below.

#### Constraints on annual reproductive success

Two types of constraint on annual reproductive success can cause the variance in offspring number in many species to be less than the Poisson expectation that 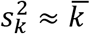. First, if maximum clutch or litter size is constrained to a small integer, that will limit the ability of some individuals to dominate reproduction, with the result that 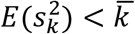 (Waples and Antao 2014). This constraint takes an extreme form in species for which females (with perhaps rare exceptions) can only produce 0 or 1 offspring in a single season; in that case, it is easy to show that 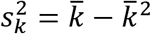, so variance in annual offspring number is always underdispersed.

The second type of constraint narrows the distribution of offspring number for one reproductive event—as commonly observed for clutch size in birds, for example (Clutton-Brock 1988). Kendall and Wittmann (2010) suggested that this phenomenon could be explained if energetic constraints and physiological feedbacks cause the instantaneous probability of laying another egg (or producing another pup) to decline with the number of eggs/pups already in the nest/litter. They proposed that the resulting distribution of clutch/litter sizes could be modeled using the generalized Poisson distribution, which has two parameters (λ_1_ and λ_2_) rather than the single parameter (λ) that characterizes the standard Poisson. The generalized Poisson distribution has a mean of 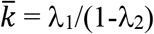 and a variance of 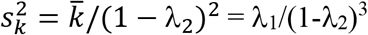, with λ_2_ constrained to be < 1 (Consul and Jain 1973). The term 1/(1-λ_2_) plays the role of a dispersion factor (Harris et al. 2012) that determines the degree of deviation from the standard Poisson: the variance is underdispersed if λ_2_ is negative, overdispersed if λ_2_ is positive, and reduces to the standard Poisson if λ_2_ = 0. Empirical data show that as maximum clutch size becomes smaller the mode approaches the maximum (Kendall and Wittmann 2010), and that pattern is reflected in the generalized Poisson distribution as λ_2_ becomes more negative. Kendall and Wittmann (2010) analyzed published data for clutch and litter size distributions for over 150 populations of terrestrial vertebrates (birds, mammals, and reptiles). They found that (1) the generalized Poisson provided a good statistical fit to 88% of the populations, much more than did other models, and (2) in all cases the λ_2_ parameter was negative, indicating 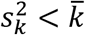.

For the analysis of clutch or litter size data for species like this, applying the *Δ_I_* adjustment proposed by Waples (2020) will overcompensate for random demographic stochasticity, leading to consistently negative estimates of *Δ_I_*. In theory, this problem has a simple solution. Let *ϕ* be the ratio of variance to mean offspring number 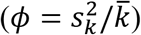 under the generalized Poisson model that allows for underdispersion. Then, the expected value of Crow’s *I* that accounts for random demographic variation in clutch or litter size is 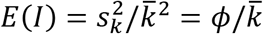. Subtracting the quantity 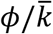 rather than 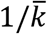 therefore will, on average, produce a generalized version of *Δ_I_* that is 0 when nothing but random demographic stochasticity is involved. We illustrate this with empirical clutch size data for the great tit (*Parus major*) from the Netherlands (Figure 4). Here, variance in clutch size (6.72) was 22% lower than the mean (8.65), so *ϕ* = 0.78—a typical result for this species. Therefore, the generalized *Δ_I_* adjustment is to subtract 0.78/8.65 = 0.090 from the raw value, rather than 1/8.65 = 0.116.

To evaluate this new adjustment, we used the above equations to solve for the two generalized Poisson parameters, producing this result:

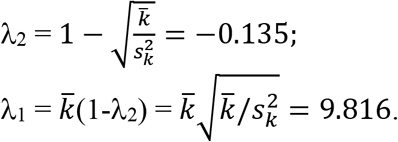

Note that λ_2_ is negative, indicating underdispersion. We then used the *rgenpois* function in R and these λ_1_ and λ_2_ values of to simulate 1000 replicate ‘populations,’ each having 217 clutch sizes (the same number for the empirical data shown in Figure 4A). For each population we calculated the mean and variance in clutch size and used these data to calculate the raw value of Crow’s *I* and the value following the generalized *Δ_I_* adjustment based on *E*(*ϕ*) = 0.78. Results (Figure 4B) show that estimates of *Δ_I_* were distributed approximately evenly around 0.

Although a generalized *Δ_I_* can be computed in this way for data from small litters or clutches, caution should be used in the interpretation. If the estimate of ϕ<1 is based on a model that quantifies the expected degree of underdispersion based on aspects of the species’ biology, then the adjusted index could be informative. However, if the estimate of ϕ<1 is based on fitting the generalized Poisson to empirical data, then the process is rather circular, as the outcome would always be *E*(generalized *Δ_I_*) = 0.

In addition, it is important to realize that an estimate close to 0 does not mean that no selection is occurring *via that fitness component*. Clutch size is known to be heritable in this (Reed et al. 2016) and other great tit populations (Santure et al 2013), meaning that additive genetic variance contributes to among-female variation in number of eggs laid, on top of demographic stochasticity. Thus, clutch size would be even more underdispersed absent this heritability, and the within-female residual variance from an ‘animal model’ (Kruuk 2004; Wilson et al. 2010) could be used to estimate the random component alone. Clutch size is itself under variable/fluctuating selection (Saether et al. 2016), and so can be considered both as a trait and a fitness component, albeit one that correlates inconsistently and sometimes not at all with annual or lifetime measures of fitness (Reed et al. 2016). Such traits/fitness components with low or highly variable ‘elasticities’ (van Tienderen 2000) are not ideal for calculating the Opportunity for Selection, as *I* or *Δ_I_* measured via these traits/fitness components might correlate poorly with *I* or *Δ_I_* measured via total fitness. The evolutionary reasons for underdispersion in clutch/litter size remain unclear, but past selection for environmental or genetic canalization might have ‘whittled away’ variation. For great tits, Mulder et al. (2016) found that within-family variance in fledgling weight (a trait correlated with clutch size) is both heritable and under stabilizing selection, indicating that adaptive evolution of trait variances as well as trait means is possible. The Opportunity for Selection is designed to provide insights into what Grafen (1988) termed ‘selection in progress’—that is, selection that occurs within the time frame encompassed by the samples being analyzed (generally within a single generation or across a generation of parent-offspring reproduction). To the extent that selection is currently operating on clutch or litter size, it likely involves selection for reaction norms that relate the mean/variance of offspring number to key environmental covariates. Evaluating this type of selection requires comparing different populations, or the same population at different times under different environmental conditions.

The reproductive constraints discussed above apply directly only to females. In the case of seasonal monogamy, similar constraints would apply indirectly to males. Furthermore, if maximum female clutch/litter size is low and the number of females a single male can access is limited, that can place an upper limit on annual reproductive success of males (as might have occurred, for example, for male black bears from Michigan; Waples et al. 2018).

#### Other factors that contribute to random stochasticity

Even in the absence of selection, other stochastic factors besides random variation in clutch/litter size can inflate the variance in offspring number and cause overdispersion compared to the Poisson expectation. Kendall and Wittmann (2010) developed a model for which the distribution of annual offspring number depends on five factors: (1) probability that the individual actually produces a clutch or a litter; (2) probability that the nest or litter survives until enumeration; (3) distribution of offspring number, contingent on (1) and (2); (4) probability that an offspring survives to independence; and (5) probability that the parent will produce one or more additional clutches/litters in the same season. So far we have only considered factor (3), but the other factors also can have a profound influence on the distribution of offspring number. All these factors can be mediated by selection but each can also have a substantial random component. Factors (1) and (2) lead to clutch/litter sizes of 0, which increase the overall variance in offspring number; these zeros are rarely included in empirical data for clutch/litter size, but it essential to consider them in an overall assessment of annual reproductive success. When variance in clutch/litter size is underdispersed, random mortality until a later life stage (factor (4)) will increase the variance-to-mean ratio (*ϕ*) to approach the Poisson expectation of 1 (Waples 2020). When it is important, factor (5) generally leads to a bimodal or multimodal distribution of offspring number, which also inflates *ϕ*.

When all these factors are considered together, the variance in annual reproductive success will often be greater than the mean, even when clutch/litter size is underdispersed. Data for the Dutch great tits, which only considered factors (3) and (5), illustrate this phenomenon. Whereas *ϕ* was <1 for clutch size, the variance of total eggs produced per year was more than twice the mean when production of multiple clutches by some birds was accounted for (Figure 4A). The generalized version of *Δ_I_* is better suited to analysis of more comprehensive data like this. If researchers can use a model like that of Kendall and Wittmann (2010), together with their knowledge of the species’ biology, to produce a comprehensive estimate of *ϕ* that accounts for as much of the random demographic stochasticity as possible (but does not soak up variation due to heritable traits affecting fitness), then the generalized 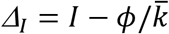 will provide a more robust estimate of the opportunity for selection.

#### Lifetime reproductive success

The cohorts that will be tracked for *LRS* include *N_z_* = *N_0_l_z_* individuals. As with the discrete generation and annual analyses, we separately consider two subsets of the cohort: those that survive to age at maturity, and those that don’t. The latter set all have 0 *LRS*, and their effect is to inflate the variance of *LRS* compared the expectation under the null model. The relative size of these two subsets of the cohort is *N_α_* = *N_0_l_α_* individuals that survive to sexual maturity and *N_die_* = *N_z_* - *N_α_* individuals that do not survive.

The numbers-that-die-at-age are given by *D_x_* = *N_x_* - *N*_*x*+*1*_, with *N*_*ω*+*1*_ = 0 because it is assumed all individuals die after reaching the maximum age (and, perhaps, producing offspring before dying). Focusing on the subset of *N_α_* individuals in the initial cohort that survived to age at maturity, the number of possible ages at death is the same as the number of ages in the adult lifespan: *AL* = ω – α + 1. The advantage of grouping individuals by age at death is that they all have the same number of seasons in which to reproduce, which simplifies calculation of the mean and variance of *LRS*. It is convenient to renumber the ages at death (*q*) as *q* = 1… *AL* (so *q* = 1 denotes individuals that survived to age at maturity but died before reaching age α + 1). With this re-indexing, mean age at death is then 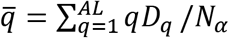.

Let *N_Off•_* be the total number of sampled offspring that are matched to the *N_α_* adults in a cohort for computation of *LRS*. Therefore, the sample mean *LRS* for the subset of the cohort that survived to age at maturity is 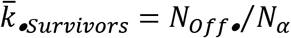, and the corresponding sample mean for the full cohort enumerated at age *z* is 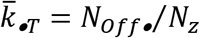. For the null model, we also need to know the expected number of offspring (*E*(*k*)) a surviving parent will produce in a single season, because all the variance components are a function of *E*(*k*). The total number of reproductive events (mature individuals alive at specific ages) for the cohort is 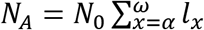, so *E*(*k*) = *N_Off•_*/*N_A_*. It follows that the expected *LRS* for individuals that die at age *q* is given by *E*(*LRS_q_*) = *qE*(*k*) and the overall mean *LRS* is 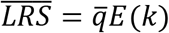.

Let 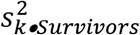 be the sample estimate of the variance in *LRS* for the subset of the cohort that survived to age α and let 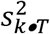 be the corresponding estimate for the entire cohort. Using the approach outlined above, together with the random expectation for 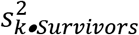 obtained by Waples (2022), we can express 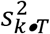 as a function of 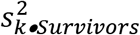 and the fractions of the cohort that did and did not survive to adulthood (see Supporting Information for details):

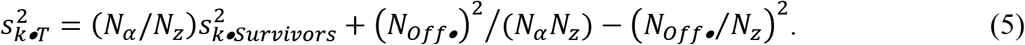

This leads to 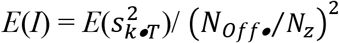

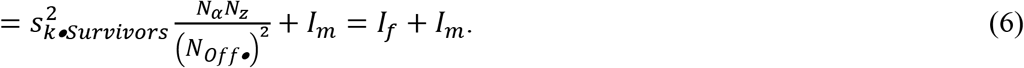

Therefore, for *LRS* as well as for annual and discrete-generation reproduction, the mortality component of *I* can be attributed to defining the cohort to include individuals that will not survive to age at maturity and hence are guaranteed to have 0 *LRS*. The remaining fraction of the overall Opportunity for Selection therefore can be attributed to *I_f_*, provided the definition of the latter is flexible enough to allow for differential lifetime fertility arising from differences in longevity.

We now focus on variance in *LRS* for individuals that do survive to adulthood, in which case the term (*N_α_*/*N_z_*) that accounts for premature mortality can be ignored. Waples (2022) derived the random *LRS* expectation for 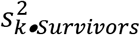. To a very good approximation, this can be simplified to (see Supporting Information for details):

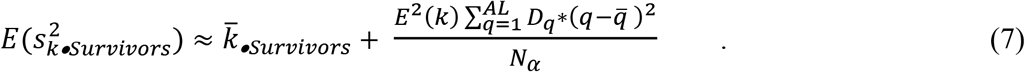

The first term in Equation 7 is the within-treatment component to variance in *LRS*, which accounts for random differences in offspring number among individuals that die at the same age. The second term accounts for random variation in lifespan (longevity). Simulation results (Figure 5) confirm the accuracy of the approximation in Equation 7.

It follows that for Crow’s index, 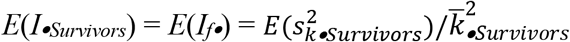

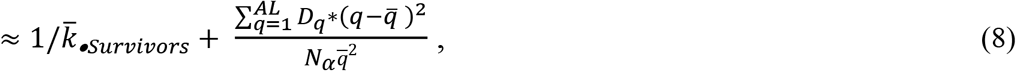

with, again, details available in Supporting Information. Equation 8 shows that under the null model for *LRS*, the expected value of the component of the Opportunity for Selection related to reproduction (*I_f•_*) is the inverse of the mean lifetime offspring number, plus a term (which does not depend on mean *LRS*) that accounts for random variation in longevity (Figure 5). This suggests an adjusted value of Δ_*I*_ for application to *LRS* data: Δ_*I•*_ = *I_f•_*– *E*(*I_f•Random_*), where *E*(*I_f•Random_*) is given by Equation 8. If population dynamics follow the null model, Δ_*I•*_ (like Δ_*I*_) has expectation 0 and does not depend on mean offspring number.

## DISCUSSION

Analytical and numerical results presented here are congruent, and they lead to the following conclusions.

### Crow’s method for partitioning the Opportunity for Selection into components related to mortality (*I_m_*) and fertility (*I_f_*) is appropriate for evaluating null models

This framework produces meaningful results for species with discrete generations, and for both annual and lifetime reproduction in age-structured species. Furthermore, this partitioning generally will be a necessary precursor to any meaningful analyses of selection.

Reproductive success data that includes both premature mortality and variation in offspring number among survivors confounds viability selection and fertility selection, making effective inference difficult or impossible.

### *I_m_* is independent of, and *I_f_* is inversely related to, mean offspring number

This result is evident from Figures 3 and 5 and applies to all three life history scenarios.

### Computing the Opportunity for Selection across an entire life cycle involves inherent tradeoffs

Total fitness of an individual can be defined as the number of offspring it produces at the start of the next generation (Walsh and Lynch 2018). To be informative with respect to total fitness, the Opportunity for Selection should be computed across an entire life cycle—otherwise, one is only studying fitness components. Quantifying offspring number in terms of zygotes producing zygotes has the advantage that it covers an entire life cycle and restricts the analysis to a single generation, making it suitable for quantitative genetic analysis (Arnold 1985; Cheverud & Moore 1994). However, unless newborns are sexually mature, zygote-to-zygote analyses will always involve both *I_m_* and *I_f_* components, which complicates interpretation unless these components can be analyzed separately.

Because tracking survival and reproduction by an entire cohort of zygotes can be logistically challenging, reproductive success in the wild is commonly measured in terms of offspring per adult, in which case production of adults by adults represents a full life cycle. Interpretation of adult-adult data in terms of the Opportunity for Selection presents two challenges. First, although the *I_m_* component of *I* disappears because there is no premature mortality, offspring mortality is subsumed into the *I_f_* component; therefore, an overall *I* value for adult-adult data reflects effects of both fertility and mortality, just as it does for zygote-zygote data. Second, the Opportunity for Selection for adult-adult data is a mixed fitness measure that applies to both parental and offspring generations, which presents well-known problems with respect to interpretation (Prout 1969; Grafen 1988; Thomson & Hadfield 2017; Walsh and Lynch 2018).

Two important tradeoffs involving the Opportunity for Selection can therefore be summarized as follows. (1) Computing the index across an entire life cycle provides information relevant to total fitness, but this complicates interpretation by combining effects of both survival and reproduction. (2) Logistical challenges in collecting zygote-zygote data can be alleviated by focusing on reproduction by adults, but this produces a mixed-fitness measure that spans parental and offspring generations. Regardless how it is measured, the Opportunity for Selection across an entire life cycle can provide useful insights regarding total fitness, but interpretation will be challenging unless the fitness components can be analyzed separately.

### The appropriate null models differ substantially for *I_m_* and *I_f_* components

**N**ull models for *I_f_* generally involve some form of random reproductive success. Nothing comparable is available for *I_m_*, which can be entirely attributed to random survival, entirely attributed to selective mortality, or some combination. Therefore, the raw *I_m_* cannot be adjusted by subtracting the expected contribution from random stochasticity, as Waples (2020) proposed to account for random reproductive success.

Instead, an appropriate null model for *I_m_* is that, if survival is random, one expects that the probability that an individual survives should be independent of its phenotype. This null model therefore can be tested by evaluating whether any observed covariance between survival and a phenotypic trait is too large to be explained by random sampling error.

### The *Δ_I_* null model often will be applicable to evaluation of *I_f_*

Waples (2020) proposed the *Δ_I_* approach based on analytical and numerical analyses of a discrete-generation model that did not consider premature mortality. Results presented here show that this still can appropriate null model for all three life histories, provided that the *I_f_* component can be isolated from *I_m_*. This can be accomplished by either a) restricting the pool of potential parents (for calculating means and variances in offspring number) to mature individuals, in which case the results apply directly to *I_f_*; or b) quantifying the magnitude of premature mortality, so that the ratio (1-*v*)/*v* can be used to quantify *I_m_*, allowing it to be subtracted from the raw *I* to yield an estimate of *I_f_*.

Analysis of lifetime reproductive success is more complicated, because in that case *I_f_* also includes a term for random variation in longevity (see Equation 8). However, the expected magnitude of the longevity term is easily calculated from cumulative, age-specific survivorship. After accounting for random variation in longevity, Crow’s Opportunity for Selection for *LRS* can be assessed using the standard Δ_*I*_ framework, the only difference being that the adjustment involves subtracting the inverse of mean 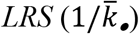 rather than the inverse of mean annual offspring number 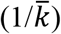.

An exception occurs for females of species for which either clutch/litter size either (1) is constrained to a small integer, or (2) has reduced variance compared to the random Poisson expectation. For these species with underdispersed variance in offspring number, using the standard adjustment proposed by Waples (2020) will lead to consistently negative *Δ_I_* values. If the expected degree of underdispersion can be quantified (in terms of the ratio ϕ), a generalized *Δ_I_* can be calculated that has an expectation of 0 when nothing but random demographic stochasticity is operating. However, as noted in Results, this approach is likely to be useful only if the expected degree of underdispersion can be quantified independently based on aspects of the species’ biology.

### A positive *Δ_I_* does not guarantee that selection is occurring

It is always important to remember that Crow’s index reflects an *opportunity* for selection but does not by itself provide direct evidence for selection. There are two major reasons for this caveat.

First, the *Δ_I_* adjustment accounts for the *expected* contribution of random (Poisson) variance in reproductive success to *I_f_*. In any real-world application, the actual magnitude of this stochastic component is a random variable that is distributed around the expected value. Therefore, a positive *Δ_I_* can occur by chance, even under the null model; conversely, *Δ_I_* can be negative if, by chance, random variation in offspring number is smaller than expected (the same phenomenon—biologically unrealistic negative estimates—can occur with other unbiased estimators, such as Weir and Cockerham’s (1984) *θ*, an analogue of *F_ST_*). Statistical significance of the *Δ_I_* index can be evaluated using the analytical expectation for the variance of 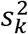 under Wright-Fisher reproduction (Waples and Faulkner 2009), or by simulations using code provided in Supporting Information.

The second major caveat is that, as discussed in Results, a variety of factors can cause non-random (family-correlated) variation in survival and/or reproduction that have nothing to do with selection. To the extent that these factors are operating, the Poisson adjustment suggested by Waples (2020) will underestimate demographic stochasticity. If the influence of these factors on the distribution of offspring number can be estimated based on what is known about the biology of the focal species, a more robust *Δ_I_* estimator can be used.

### Other factors that can be important

Our null model for *LRS* assumes that survival and reproduction are independent over time. In the real world, both positive and negative correlations are common. Persistent individual differences (some individuals being consistently above or below average at producing offspring) leads to positive temporal correlations in offspring number, increases var(*LRS*) (Lee et al. 2011), and generally would be expected to lead to a positive *Δ_I_*. Conversely, intermittent breeding generally reduces var(*LRS*) and hence *Δ_I•_*, because it limits opportunities for some individuals to consistently dominate offspring production (Waples and Antao 2014). For either scenario, comparison with a null model that assumes independence of survival and reproduction over time can provide a valuable reference point for interpreting empirical data.

Intermittent breeding also has another consequence for annual reproduction: it reduces the number of breeders and creates two classes of potential parents—those that will participate in reproduction that season, and those that won’t. The latter subset by definition all leave zero offspring, so including them in the calculation of the mean and variance in annual reproductive success has the same qualitative consequences as does premature mortality. Two general options are available to deal with this issue. First, if non-breeding adults are included in the calculations of 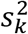 and 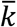, the null model could be adjusted to become a zero-inflated Poisson, as described in Supporting Information. This option only requires an estimate of the fraction of non-breeders each year. A second, simpler option is to use only breeders to compute 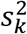 and 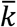, in which case the analysis can proceed as described above.

Analyses presented here have all assumed that, although age at maturity might vary among populations, within a population a single α applies to all individuals. In many real populations age at maturity varies among individuals, in which case at some ages there is a mix of mature and immature individuals. By definition all the immature individuals produce zero offspring until they mature, so for those mixed ages the consequences are similar to having a mix of breeders and non-breeders. If immature individuals can be identified, they can be excluded from the pool of potential parents used to compute 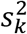 and 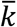; if not but the fraction mature at each age can be estimated, this information could be used to compute an analogue to *I_m_* that can be removed, so analysis can focus on the *I_f_* component.

## Acknowledgments

This manuscript benefitted considerably from discussions with Joel Pick and Michael Morrissey. The authors declare no conflict of interest.

## Data Availability

Data for *Parus major* used in Figure 4 will be archive in Dryad on acceptance.

