## Supplemental info for "Null models for the Opportunity for Selection"

### Supporting Information

#### I. Zero inflated Poisson

Consider two processes: Process A always produces zeroes, and Process B produces random Poisson variables, some of which might be zero. A mixture of a fraction $\pi$ outcomes from B and a fraction (1- $\pi$) outcomes from A has a zero-inflated Poisson distribution. If the Poisson parameter for Process B is $\lambda$, the mixed distribution has a parametric mean of $\mu = \pi\lambda$ and a parametric variance of

$$\sigma^2 = \mu + \mu^2[(1-\pi)/\pi]$$
$$= \pi\lambda + (\pi\lambda)^2[1/\pi - 1].$$

Now let $\pi = v$ = the fraction of individuals that survive from an earlier life stage to adulthood, let 1- $v$ be the fraction that don't survive, and let $\bar{k}$ = mean offspring number for the survivors be the Poisson parameter ($\lambda$). Then for the mixed distribution, the sample means and variances are given by

$$\bar{k}_T = v\bar{k} \quad \text{and}$$
$$s_T^2 = \bar{k}_T + (\bar{k}_T)^2[\tfrac{1}{v} - 1],$$

as obtained in the main text using another method.

A similar zero-inflated phenomenon can occur without mortality if there are two classes of adults: those that participate in reproduction and have a chance to produce offspring, and those that don't. Non-breeding can be a permanent condition (sterility) or a yearly option (intermittent breeding). Consider first the subset of individuals that participates in reproduction, and for this subset let the relationship between the mean and variance in offspring number be $s_{k(b)}^2 = \phi_b \bar{k}_b = \phi_b \Sigma k / n_b$, where the subscript '$b$' indicates these parameters apply to the $n_b$ breeders. Letting $SS_b$ be the sum of squared offspring number for the breeders, from the definition of a variance it follows that

$$s_{k(b)}^2 = \frac{SS_b}{n_b} - (\bar{k}_b)^2 = \phi_b \bar{k}_b = \frac{\phi_b \Sigma k}{n_b}. \qquad \text{(S1)}$$

For the breeders, Crow and Morton's (1995) 'Index of Variability' is $s_{k(b)}^2/\bar{k}_b = \phi_b$, and the fertility component of the Opportunity for Selection is

$$I_{f(b)} = \frac{s_{k(b)}^2}{\bar{k}_b^{\,2}} = \frac{\phi_b}{\bar{k}_b}, \text{ and} \qquad \text{(S2)}$$

$$\Delta_{I_{f(b)}} = \frac{\phi_b}{\bar{k}_b} - \frac{1}{\bar{k}_b} = \frac{\phi_b - 1}{\bar{k}_b}. \qquad \text{(S3)}$$

Note that $\Delta_{I_{f(b)}}$ is 0 if the breeders have Poisson variance in reproductive success.

Because non-breeders produce no offspring, they don't contribute to either $\Sigma k$ or $SS_b = \Sigma k^2$. For the following analyses, it is convenient to have an expression for $SS_b$, which is obtained by solving Equation S1 for $SS_b$:

$$SS_b = n_b\,[\phi_b \Sigma k/n_b + (\Sigma k/n_b)^2].$$

Assume now that some individuals are non-breeders, such that the total number of individuals is some multiple of the number of breeders: $n_T = Xn_b$. The new overall mean offspring number is then $\bar{k}_T = \Sigma k/n_T = \Sigma k/(Xn_b)$, and the overall variance is given by

$$s^2_{k(T)} = \frac{SS_b}{n_T} - \left(\bar{k}_T\right)^2 = \frac{n_b\,[\phi_b \Sigma k/n_b + (\Sigma k/n_b)^2]}{Xn_b} - \left(\frac{\Sigma k}{Xn_b}\right)^2$$

$$= \frac{[\phi_b \Sigma k/n_b + (\Sigma k/n_b)^2]}{X} - \left(\frac{\Sigma k}{Xn_b}\right)^2$$

$$= \frac{\phi_b \Sigma k}{Xn_b} + \frac{(\Sigma k)^2}{X(n_b)^2} - \frac{(\Sigma k)^2}{X^2(n_b)^2}$$

$$= \frac{\Sigma k}{Xn_b}\left[\phi_b + \frac{\Sigma k}{n_b} - \frac{\Sigma k}{Xn_b}\right]$$

$$= \bar{k}_T[\phi_b + \bar{k}_b - \bar{k}_T]. \qquad \text{(S4)}$$

For the overall population, therefore, the Index of Variability is

$$\phi_T = s^2_{k(T)}/\bar{k}_T = \phi_b + \bar{k}_b - \bar{k}_T,$$

and the Opportunity for Selection is

$$I_{f(T)} = \frac{s^2_{k(T)}}{\bar{k}_T^{\,2}} = \frac{\phi_T}{\bar{k}_T} = \frac{(\phi_b + \bar{k}_b)}{\bar{k}_T} - 1 = \frac{\phi_b}{\bar{k}_T} + \frac{\bar{k}_b}{\bar{k}_T} - 1$$

$$= \frac{\phi_b}{\bar{k}_T} + \frac{\frac{\Sigma k}{(n_b)}}{\frac{\Sigma k}{(Xn_b)}} - 1 = \frac{\phi_b}{\bar{k}_T} + \frac{n_T}{n_b} - 1. \qquad \text{(S5)}$$

Equation S5 shows that in this mixed population of breeders and non-breeders, the overall Opportunity for Selection has two components: the first component ($\frac{\phi_b}{\bar{k}_T}$) reflects the variance in reproductive success among the breeders, and the second component ($\frac{n_T}{n_b} - 1$) reflects the effects of zero inflation due to non-breeders. If there are no non-breeders ($n_T$=$n_b$), this second component disappears. Applying the $\Delta_I$ correction produces

$$\Delta_{I_{f(T)}} = \frac{\phi_b}{\bar{k}_T} + \frac{n_T}{n_b} - 1 - \frac{1}{\bar{k}_T} = \frac{(\phi_b - 1)}{\bar{k}_T} + \frac{n_T}{n_b} - 1.$$

In the special case where the breeders have Poisson reproductive success ($\phi_b$=1), this reduces to

$$\Delta_{I_{f(T)}} = \frac{n_T}{n_b} - 1,$$

in which case all of the overdispersion compared to the Poisson expectation arises from the fact that all non-breeders produce zero offspring.

### II. Lifetime reproductive success

Let $s^2_{k\bullet Survivors}$ be the sample estimate of the variance in *LRS* for the subset of the cohort that survived to age α and let $s^2_{k\bullet T}$ be the corresponding estimate for the entire cohort. Using the approach outlined above, we can express $s^2_{k\bullet T}$ as a function of $s^2_{k\bullet Survivors}$ (for which an analytical solution was provided by Waples 2022) and the fractions of the cohort that did and did not survive to adulthood. For the subset of survivors, under the null model

$$SS_{Adult} = N_\alpha * \left[s^2_{k\bullet Survivors} + \left[N_{Off\bullet}/N_\alpha\right]^2\right],$$

and this is also the total *SS*. Therefore,

$$s^2_{k\bullet T} = SS_{Adult}\,/N_z - \left(N_{Off\bullet}/N_z\right)^2$$
$$= (N_\alpha/N_z)\left[s^2_{k\bullet Survivors} + \left[N_{Off\bullet}/N_\alpha\right]^2\right] - \left(N_{Off\bullet}/N_z\right)^2$$
$$= (N_\alpha/N_z)s^2_{k\bullet Survivors} + (N_\alpha/N_z)\left[N_{Off\bullet}/N_\alpha\right]^2 - \left(N_{Off\bullet}/N_z\right)^2$$
$$= (N_\alpha/N_z)s^2_{k\bullet Survivors} + \left(N_{Off\bullet}\right)^2/(N_\alpha N_z) - \left(N_{Off\bullet}/N_z\right)^2.$$

This leads to $E(I) = E(s^2_{k\bullet T})/\left(N_{Off\bullet}/N_z\right)^2$

$$= \frac{(N_\alpha/N_z)s^2_{k\bullet Survivors} + \left(N_{Off\bullet}\right)^2/(N_\alpha N_z) - \left(N_{Off\bullet}/N_z\right)^2}{\left(N_{Off\bullet}/N_z\right)^2}$$
$$= s^2_{k\bullet Survivors}\frac{N_\alpha N_z}{\left(N_{Off\bullet}\right)^2} + \frac{N_z}{N_\alpha} - 1$$
$$= s^2_{k\bullet Survivors}\frac{N_\alpha N_z}{\left(N_{Off\bullet}\right)^2} + \frac{N_z - N_\alpha}{N_\alpha}$$

$$= s^2_{k\bullet Survivors}\frac{N_\alpha N_z}{\left(N_{Off\bullet}\right)^2} + I_m = I_f + I_m. \qquad \text{(S6)}$$

From Waples (2022), using current notation:

$$E(s^2_{k\bullet Survivors}) = [E(k)/N_\alpha]\sum_{q=1}^{AL} q\left(D_q - 1\right) \quad \text{(within treatment)}$$
$$+[E(k)/N_\alpha]\sum_{q=1}^{AL} q\,\frac{(D_q-1)}{D_q} \quad \text{(among ages)}$$
$$+\frac{\sum_{q=1}^{AL} D_q * (q*E(k) - \bar{k}_{\bullet Survivors})^2}{N_\alpha} \quad \text{(longevity)}$$

Note that $q$ = age at death, $\bar{q} = \sum_{q=1}^{AL} qD_q\,/N_\alpha$, and $\bar{k}_{\bullet Survivors} = E(k)\bar{q}$. A very good approximation to the above, which considerably simplifies interpretation, can be obtained by 1) ignoring the 'among ages' component, which generally will be a small fraction of the total, and 2) ignoring the '-1' term in the 'within treatment' component. These two adjustments have opposite effects and largely cancel out. With these changes:

$$E(s^2_{k\bullet Survivors-Within}) \approx E(k)\sum_{q=1}^{AL} qD_q/N_\alpha = E(k)\bar{q} = \bar{k}_{\bullet Survivors}$$

and

$$E(s^2_{k\bullet Survivors-Longevity}) = \frac{\sum_{q=1}^{AL} D_q*(qE(k)-\bar{k}_{\bullet Survivors})^2}{N_\alpha}$$

$$= (\frac{1}{N_\alpha})\sum_{q=1}^{AL} D_q(qE(k) - E(k)\bar{q}\ )^2 = \frac{E^2(k)\sum_{q=1}^{AL} D_q*(q-\bar{q}\ )^2}{N_\alpha}\ .$$

So total $E(s^2_{k\bullet Survivors}) \approx \bar{k}_{\bullet Survivors} + \frac{E^2(k)\sum_{q=1}^{AL} D_q*(q-\bar{q}\ )^2}{N_\alpha}$ . (S7)

It follows that $E(I_{\bullet Survivors}) = E(I_{f\bullet}) = E(s^2_{k\bullet Survivors})/\bar{k}^2_{\bullet Survivors}$

$$\approx 1/\bar{k}_{\bullet Survivors} + \frac{E^2(k)\sum_{q=1}^{AL} D_q*(q-\bar{q}\ )^2}{N_\alpha \bar{k}^2_{\bullet Survivors}}.$$

Now the lifetime mean offspring number is proportional to the mean annual offspring number as follows: $\bar{k}_{\bullet Survivors} = \bar{q}E(k)$, with the proportional constant being mean age at death. Making this substitution produces

$$E(I_{f\bullet}) = 1/\bar{k}_{\bullet Survivors} + \frac{E^2(k)\sum_{q=1}^{AL} D_q*(q-\bar{q}\ )^2}{N_\alpha E^2(k)\bar{q}^2}$$

$$= 1/\bar{k}_{\bullet Survivors} + \frac{\sum_{q=1}^{AL} D_q*(q-\bar{q}\ )^2}{N_\alpha \bar{q}^2}\ , \qquad \text{(S8)}$$

where the first term reflects Poisson variance in lifetime reproductive success among individuals that die at the same age, and the second term reflects differential mean *LRS* for individuals that die at different ages.

Note that in Equation S7 for $E(s^2_{k\bullet Survivors})$, the second (longevity) term is proportional to the square of mean annual offspring number. Converting this to $E(I_{\bullet Survivors})$ involves division by the square of mean lifetime offspring number, which cancels that effect. As a consequence, the longevity component of $E(I_{\bullet Survivors})$ is independent of both $E(k)$ and $\bar{k}_{\bullet Survivors}$.

### III. Computer code

#### Code for the analyses in Figure 3

```
omega = 9
alpha = 3
AL = omega-alpha+1
z = 1
Ages = 0:omega
survival = 0.7
N1 = 800
NReps = 100
multiple = c(0.5,1,2,5,10)
kbar = matrix(NA,NReps,length(multiple))  ## actual mean fecundity of adults
kbarT = kbar  ## mean across all sampled parents
Vk = kbar
VkT = kbar
If = kbar
IfT = kbar

lx = 1:(omega+1)
for (j in 2:(omega+1)) {lx[j] = lx[j-1]*survival}
NParents = 1:(omega+1)


for (M in 1:length(multiple)) {
lambda = multiple[M]


for (R in 1:NReps)  {

NParents[1] = N1
for (j in 2:omega)  { NParents[j] = rbinom(1,NParents[j-1],survival) }
NAdults = sum(NParents[(alpha+1):(omega+1)])
TotN = sum(NParents[(z+1):(omega+1)])
Im = (TotN-NAdults)/NAdults

Offspring = rpois(NAdults,lambda)
kbar[R,M] = mean(Offspring)
Vk[R,M] = var(Offspring)
kbarT[R,M] = sum(Offspring)/TotN
VkT[R,M] = sum(Offspring^2)/TotN - kbarT[R,M]^2
If[R,M] = 1/kbar[R,M]
IfT[R,M] = VkT[R,M]/(kbarT[R,M])^2 - Im

}  ## end for R
}  ## end for M

EVk = kbar
EVkT = kbarT + kbarT^2*(TotN-NAdults)/NAdults
EIf = 1/kbar
EIfT = 1/kbarT

###comparing observed and expected means for different values of kbar
colMeans(kbar)/multiple
colMeans(Vk)/colMeans(EVk)
```

```
colMeans(VkT)/colMeans(EVkT)
colMeans(If)/colMeans(EIf)
colMeans(IfT)/colMeans(EIfT)

CIS = array(data = NA, dim = c(3,5,length(multiple)),dimnames =
list(c("low","median","hi"),c("kbar","Vk","VkT","If","IfT"),multiple))

for (M in 1:length(multiple)) {
  CIS[,1,M] = quantile(kbar[,M],probs = c(0.025, 0.5, 0.975))
   CIS[,2,M] = quantile(Vk[,M],probs = c(0.025, 0.5, 0.975))
    CIS[,3,M] = quantile(VkT[,M],probs = c(0.025, 0.5, 0.975))
     CIS[,4,M] = quantile(If[,M],probs = c(0.025, 0.5, 0.975))
      CIS[,5,M] = quantile(IfT[,M],probs = c(0.025, 0.5, 0.975))

  }  ## end for M

### The array CIs gives more information about the distribution of offspring number
```

### Code for the analyses in Figure 5

```
omega = 9
alpha = 3
AL = omega-alpha+1
z = 1
AL = omega - alpha + 1  ## adult lifespan
Ages = 1:AL
survival = 0.7
N1 = 800
Nx = 1:(omega+1)
NReps = 1000
multiple = c(0.5,1,2,5,10)
MeanLRS = 2*multiple

TotalSS = matrix(NA,NReps,length(multiple))
Vk = TotalSS
BetweenSS = TotalSS
WithinSS = TotalSS
LongevitySS = TotalSS
Crow = TotalSS
within = 1:AL
between = 1:AL
within2 = 1:AL
between2 = 1:AL
long = 1:AL
ADkbar = matrix(NA,NReps,AL)
ADVk = ADkbar
ADkbar2 = ADkbar
ADVk2 = ADkbar

for (M in 1:length(multiple))  {

for (R in 1:NReps)  {

### get realized age structure for a cohort, allowing for random survival
```

```
Died = rep(NA,AL)
Nx[1] = N1
for (j in 2:(omega+1))  { Nx[j] = rbinom(1,Nx[j-1],survival) }
for (j in 1:(AL-1))  { Died[j] = Nx[j+alpha] - Nx[j+alpha+1] }
Died[AL] = Nx[omega+1]

Nalpha = Nx[alpha+1] ## number of the cohort that reached adult
NAdults = sum(Nx[(alpha+1):(omega+1)])
TotN = Nx[z+1] ## total number in the cohort
Im = (TotN-NAdults)/NAdults
TotOff = MeanLRS[M]*Nalpha
kbar = TotOff/NAdults

Pdata = matrix(NA,Nx[alpha+1],AL)
IDs = 1:Nalpha
Pdata = cbind(IDs,Pdata)

## Model reproduction at age alpha
Offspring = rpois(Nalpha,kbar)
Pdata[,2] = Offspring

Survivors = list()
for (j in 1:AL)  {
 Survivors[[j]] = 1:Nx[alpha+j]  }

Survivors[[1]] = IDs
## get survivors to subsequent ages , and their offspring production
for (j in 2:AL)  {
  NewN = Nx[alpha+j]
  Survivors[[j]] = sample(Survivors[[j-1]],NewN,replace=F)
  NewOffspring = rpois(NewN,kbar)
    for (k in 1:NewN) {
      row = Survivors[[j]][k]
      Pdata[row,j+1] = NewOffspring[k] }  ## end for k
    } ## end for j

V = 1:AL
for (j in 1:AL)  {
V[j] = var(Pdata[,j+1],na.rm=T)
}  # end for j
V/kbar

LRO = rowSums(Pdata[,-1],na.rm=T)
Vk[R,M] = var(LRO)
AllK = sum(LRO)
BigMean = AllK/Nalpha
Crow[R,M] = Vk[R,M]/BigMean^2
TotalSS[R,M] = (Nalpha)*Vk[R,M]  ## total sum of squares

                ##Get age at death
                PD = Pdata[,-1]
                AgeDeath = rep(AL,Nalpha)
                for (k in 1:Nalpha)  {
```

```
            X = which(is.na(PD[k,]))
            if(length(X)>0)  {AgeDeath[k] = X[1]-1}
            }  # end for k
            table(AgeDeath)

### analyze by age at death
for (j in 1:AL)  {
  bit = subset(Pdata,AgeDeath == j)
  bit = bit[,-1]
  X = nrow(bit)
  LRObit = rowSums(bit,na.rm=T)
  ADkbar[R,j] = mean(LRObit)
  dev = LRObit-mean(LRObit)
  within[j] = sum(dev^2)
  ADVk[R,j] = within[j]/X
  between[j] = X*(mean(LRObit)-BigMean)^2
}  ## end for j

WithinSS1 = sum(within)
BetweenSS1 = sum(between)

##Repeat after removing within-age and between-age effects
Pdata2 = Pdata[,-1]
Not = sum(is.na(Pdata2))
Used = AL*Nalpha-Not
Bigkbar = AllK/Used
for (k in 1:Nalpha)  {
for (j in 1:AL)  {
 if(is.na(Pdata2[k,j])== FALSE) {
 Pdata2[k,j] = Bigkbar }  ## end if
 }} ## end for k,j

for (j in 1:AL)  {
 bit2 = subset(Pdata2,AgeDeath == j)
 X = nrow(bit2)
 LRObit = rowSums(bit2,na.rm=T)
 ADkbar2[R,j] = mean(LRObit)
 dev = LRObit-mean(LRObit)
 within2[j] = sum(dev^2)
 ADVk2[R,j] = within2[j]/X
 between2[j] = X*(mean(LRObit)-BigMean)^2
}  ## end for j

WithinSS2 = sum(within2)
BetweenSS2 = sum(between2)

WithinSS[R,M] = WithinSS1
BetweenSS[R,M] = BetweenSS1
LongevitySS[R,M] = BetweenSS2

}  # end for R

}  # end for M
```

```
### get general expectations for variance components based on parametric vital rates
ENx = 1:(omega+1)
ENx[1] = N1
for (j in 2:(omega+1)) { ENx[j] = round(ENx[j-1]*survival)}
ENalpha = ENx[alpha+1]
ENAdults = sum(ENx[(alpha+1):(omega+1)])
EDied = rep(NA,AL)
for (j in 1:(AL-1))  { EDied[j] = ENx[j+alpha] - ENx[j+alpha+1] }
EDied[AL] = ENx[omega+1]

ELRObar = MeanLRS
Ekbar = MeanLRS*ENalpha/ENAdults
ELRO = matrix(NA,length(multiple),AL)
EWRandom = ELRO
ELongRandom = ELRO

for (M in 1:length(multiple))  {
for (j in 1:AL)  {
 ELRO[M,j] = j*Ekbar[M]
 EWRandom[M,j] = (EDied[j])*ELRO[M,j]
 ELongRandom[M,j] =  EDied[j]*(j*Ekbar[M]-ELRObar[M])^2
}}  # end for M,j

  ESSWRandom = rowSums(EWRandom)
  ESSLongRandom = rowSums(ELongRandom)
  ESSTRandom = ESSWRandom+ESSLongRandom

EVkRandom = ESSTRandom/ENalpha
ECrow = EVkRandom/ELRObar^2

EVkRandom  ## Expected var(LRS)
colMeans(Vk)  ## actual var(LRS)
ECrow
colMeans(Crow) ## actual Crow's If
```
